# Tfh dysfunction is associated with poor responses to HBV vaccination

**DOI:** 10.64898/2026.07.30.741822

**Authors:** Han M. Chen, Elizabeth Severa, Xiaomin Yao, Tijaana Williams, Riddhimaa Sinha, Clarice Monteiro, Michael Tuen, Sam Barnett Dubensky, Sergei B. Koralov, Carla R. Nowosad, Derek A. Oldridge, Laura A. Vella, Anoma Nellore, Ramin Sedaghat Herati

## Abstract

The primary goal of vaccination is to induce durable protection through generating antigen-specific humoral immunity and cellular immune memory. Yet the same vaccine may elicit protection in some and not in others. To characterize the molecular and cellular events that underlie differential vaccine responsiveness, we leveraged the known, pronounced variability in vaccine antigen-specific antibody concentration and durability observed after Hepatitis B virus (HBV) immunization. We longitudinally profiled 101 healthy adults undergoing either *de novo* (n=59) or booster (n=42) HBV immunizations. We then stratified participants into High (≥100 mIU/mL) and Low (<100 mIU/mL) Responder groups based on end-of-study titers. From this cohort, we obtained core needle biopsies (CNBx) of vaccine-draining lymph nodes from 10 participants (High Responder n=6, Low Responder n=4), 10-21 days post-final vaccination and performed single-cell RNA sequencing with paired antigen receptor sequencing. Lymph node BCR repertoire analysis revealed convergent, semi-public clonotypes across unrelated participants with elevated somatic hypermutation. Transcriptomic profiling of GC-B cells uncovered functional divergence between response groups, with High Responders showing robust activation across both dark zone (DZ) and light zone (LZ) compartments and enrichment of MYC and mTORC1 signaling. This divergence in B cell responses corresponded to notable differences in the T cell compartment. High Responders harbored a key Tfh subset marked by increased expression of CXCL13, ICOS, and GNG4, supportive of their GC localization and capacity to provide B cell help. Low Responders demonstrated a divergent program in which their GNG4+Tfh displayed aberrant inflammatory transcriptional activity despite expressing coordinated costimulatory genes necessary to provide help to LZ GC-B cells. This inflammatory signaling was associated with reduced specificity of *BCL6* regulon activity despite maintained expression of canonical Tfh genes. Our data suggest that variable HBV vaccine responsiveness was associated with transcriptional aberrations in the GNG4+Tfh subset, offering a molecular framework for understanding poor vaccine responses.

**One Sentence Summary:** Lymph node Tfh dysfunction is associated with poor vaccine responses to hepatitis B virus vaccination.

## INTRODUCTION

Although Hepatitis B virus (HBV) vaccination is broadly recommended, non-responsiveness to the HBV vaccine poses a significant public health challenge, leaving millions vulnerable to chronic infection, cirrhosis, and hepatocellular carcinoma. Despite multiple doses, 5-10% of adults fail to achieve seroprotective titers (*1*, *2*). Chronic inflammatory states, including aging (*3*), obesity (*4*, *5*), cigarette smoking (*6*), and renal disease (*7*) are associated with diminished HBV vaccine responses; however, these factors do not fully account for the heterogeneity in responses. Understanding the basis of this heterogeneity therefore requires characterization of underlying cellular processes that shape humoral immunity at the site of the response.

The overarching goal of HBV vaccination is to establish durable, antigen-specific immune memory. Optimal humoral responses depend on germinal centers (GCs), specialized structures critical for affinity maturation, clonal selection, and differentiation of memory B cells (MBCs) and long lived plasma cells (LLPCs) (*8–12*). Within GCs, T follicular helper (Tfh) cells provide essential help to cognate B cells, driving somatic hypermutation and clonal selection (*12–14*). Tfh subsets encompassing distinct functional states have been associated with vaccine-induced immune responses (*14–17*). Yet, vaccination elicits highly variable humoral responses (*18*, *19*), and the cellular and molecular programs underlying these variable responses in humans remain unclear.

Despite rich mechanistic insight from murine models, our understanding of human GC dynamics in the context of vaccination remains nascent. In humans, circulating Tfh (cTfh) and antibody secreting cells (ASCs) from peripheral blood following vaccination serve as accessible correlates of GC dynamics (*20–26*). While cTfh cells and circulating ASCs have provided valuable peripheral correlates, a major methodological gap persists in human immunology: peripheral blood proxies do not fully capture the complex cellular dynamics and localized microenvironments within secondary lymphoid organs where GC reactions take place. To close this gap, recent studies deploying fine-needle aspiration (FNA) of vaccine-draining lymph nodes have demonstrated coordinated peripheral and GC responses in influenza (*27–29*) and SARS-CoV2 mRNA vaccinations longitudinally (*30–32*), as well as identifying GC dysfunction as a potential cause for impaired responses in immunocompromised kidney transplant recipients (*33*). Together these studies establish lymph node sampling as a valuable tool to study GC responses in humans, offering a window into key processes necessary for immunity. However, we still lack understanding of why humoral responses vary between individuals, particularly with respect to molecular drivers of dysfunction.

To better characterize the link between GC responses and variability in humoral responses, we recruited ambulatory adults into a longitudinal study of HBV vaccination (NCT04674462), comprising HBV-vaccine-naive adults who received *de novo* vaccine series (n=59) and adults with prior HBV vaccination history who received a single booster (n=42). Participants were stratified into High Responders and Low Responders based on final anti-HBV antibody titers. In addition, we performed an ultrasound-guided core needle biopsy (CNBx) of enlarged ipsilateral axillary lymph nodes in ten individuals within 10-21 days following the final vaccination. Immunofluorescence confirmed the presence of GCs. Single-cell B cell receptor (BCR) analysis identified convergent GC-B cell responses including public clonotypes across participants. High Responders exhibited hallmarks of robust GC responses relative to Low Responders, including highly mutated repertoire and proliferative capability in GC-B cells. Moreover, in High Responders, GC-Tfh cells maintained canonical costimulatory programs and robust T-B cell interaction networks required for B cell survival and affinity maturation, whereas Low Responders had fewer GC-Tfh as well as dysfunctional gene regulatory networks despite greater expression of some canonical Tfh genes. Together, these findings emphasize the requirement for the coordinated engagement of GC-B and Tfh cells for productive responses and reinforce the need for further studies of Tfh to better understand suboptimal vaccine responses.

## RESULTS

### Heterogeneous humoral responses following Hepatitis B vaccination in adults

To interrogate the cellular and molecular mechanisms driving differential responsiveness to HBV vaccination, we prospectively enrolled 101 ambulatory adults in two main cohorts: a *de novo* vaccination cohort of HBV-naive adults and a booster cohort of adults with reported prior HBV immunization (NCT04674462; **Fig. 1A and S1A**). The median age of participants in the study was 54 (**Table S1**). Individuals without prior HBV vaccination history (HBVnaive) were randomized in a 1:4 ratio to receive either two doses of CpG-adjuvanted (HBVnaive-CpG, n=13) or three doses of alum-adjuvanted vaccine (HBVnaive-Alum, n=46), whereas those with prior HBV vaccination (HBVimmune, n=42) received a single alum-adjuvanted dose (**Fig. 1A and S1A**). As per package insert guidelines, HBVnaive-Alum participants received vaccinations at approximately 0, 4 and 24 weeks after study entry, whereas HBVnaive-CpG received vaccinations at 0 and 4 weeks. For all three study arms, we collected blood at the time of vaccination (w0), one week after each vaccination (w1) and four weeks after vaccination (w4), facilitating longitudinal analysis of antibody kinetics across successive vaccinations. In total, we collected 531 blood samples from the 101 participants. Additionally, participants who consented underwent screening for enlarged, ipsilateral axillary lymph nodes amenable to core needle biopsy (CNBx) 10 to 21 days after final vaccination, in order to capture early GC dynamics following immunization (*31*, *34–36*).

**Figure 1.**
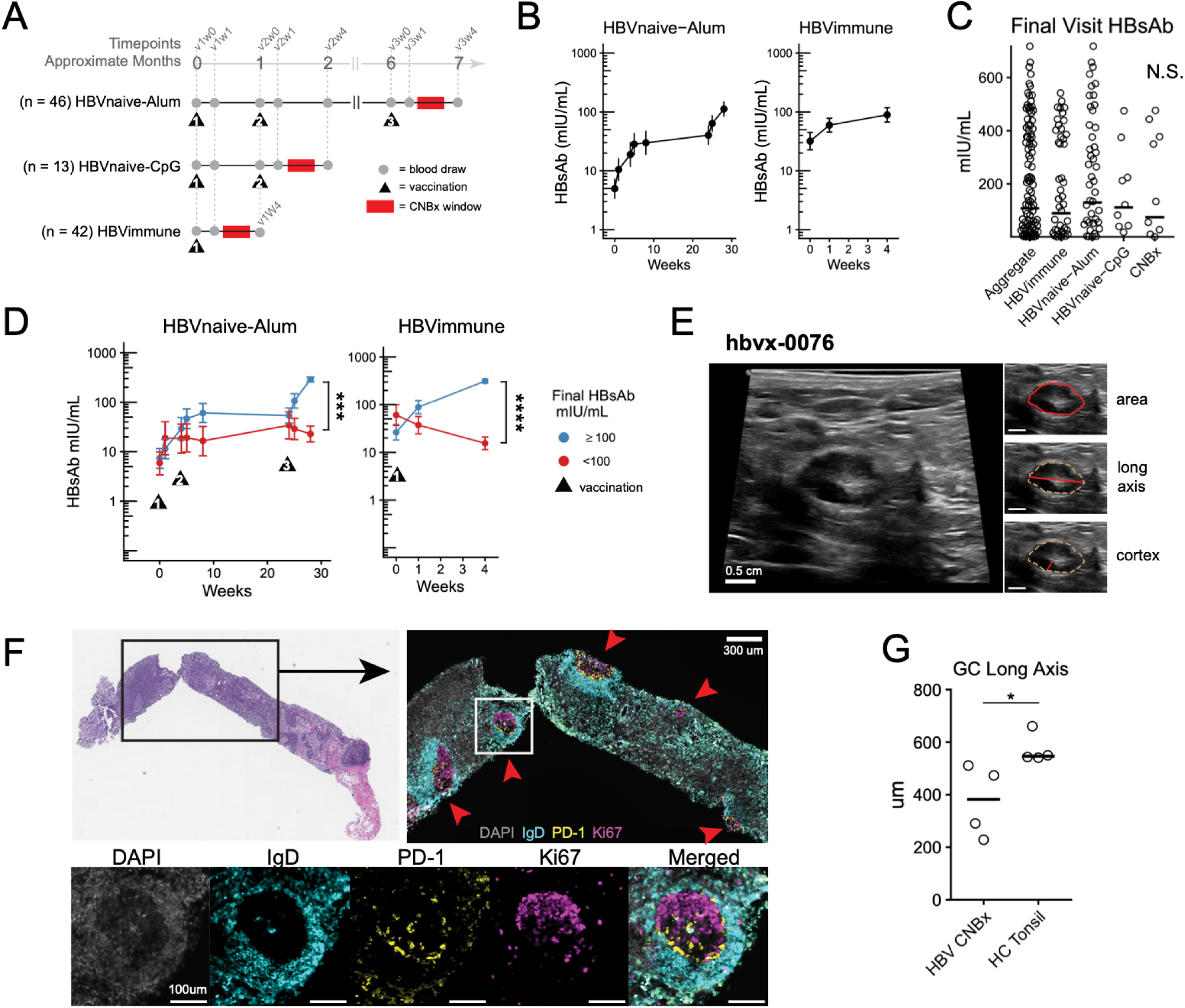
Heterogeneous humoral responses following Hepatitis B vaccination in adults. **A.** Timeline of the clinical study outlining vaccination schedules (black triangles, numbered by dose), blood draw timepoints (gray circles), and core needle biopsy (CNBx) window (red bars) across three cohorts: HBVnaive-Alum (n=46), HBVnaive-CpG (n=13), and HBVimmune-Alum (n=42). Approximate months since initial visit are shown on upper x-axis, with sample timepoints labeled (e.g. v1w0 = visit 1, week 0). **B.** Longitudinal total HBsAb titers (mIU/mL) over time for the HBVnaive-Alum and HBVimmune cohorts, illustrating the overall rise in antibody titer following each vaccine dose. **C.** Distribution of HBsAb titers at the final study visit, shown for the aggregate cohort, each individual study arm, and the subset of participants who underwent CNBx. Horizontal lines indicate geometric means. Final titers were similar across arms (N.S.), and the CNBx subcohort HBsAb distribution was representative of the larger cohort. **D.** Longitudinal HBsAb kinetics stratified by final visit HBsAb titer, with participants categorized into High Responders (≥100 mIU/mL, blue) and Low Responders (<100 mIU/mL, red). Data show geometric mean HBsAb titers ± SEM over time, with black triangles marking vaccination timepoints and number indicating dose. P-values (HBVnaive-Alum P=3.12e-4; HBVimmune P=2.48e-9) represent the group-by-time interaction from linear mixed effects models fit to log-transformed HBsAb titers, with Satterthwaite-approximated degrees of freedom (lme4/lmerTest). **E.** Representative ultrasound imaging (participant hbvx-0076) of an enlarged ipsilateral axillary lymph node following vaccination. Insets details denote cross-sectional area (red outline), long axis and cortical thickness (red segments). Scale bar = 0.5 cm. **F.** Histologic confirmation of GC activity within a biopsied lymph node. Top left: low-power brightfield overview of a hematoxylin and eosin (H&E)-stained biopsy core; boxed area indicates the region magnified at top right. Top right: low-power immunofluorescence for DAPI (gray), IgD (cyan), PD-1 (yellow), and Ki67 (magenta); red arrowheads denote follicular structures containing GCs. Scale bar = 300 µm. Bottom: high-magnification view of a single GC. Scale bar = 100 µm. **G.** Quantitative comparison of GC long axis diameters (µm) between healthy control tonsil tissue (HC Tonsil) and vaccine draining lymph node biopsies (HBV CNBx). *P < 0.05. N.S., non-significant.

To track humoral responses to the vaccine immunogen, HBV surface antigen (HBsAg), we quantified systemic HBsAg-specific antibodies (HBsAb) by ELISA. In each of the three study arms, we observed overall HBsAb increase following vaccination, with a geometric mean increase relative to baseline of 2.8-fold for the HBVimmune group, 16-fold for HBVnaive-Alum and 6.9-fold for HBVnaive-CpG (**Fig. 1B and S1B**). Similar distribution of end-of-study HBsAb was observed between the HBVimmune and HBVnaive-Alum groups (P>0.05, n=42 vs n=45, one-way ANOVA with Tukey’s post-test; **Fig. 1C**). Aggregate end of study geometric mean HBsAb was 97 mIU/mL, with group-level means of 89 mIU/mL (HBVimmune), 101 mIU/mL (HBVnaive-Alum), and 122 mIU/mL (HBVnaive-CpG). We therefore stratified participants into High and Low Responders (≥100 vs. <100 mIU/mL) to examine factors that distinguished the response groups. Although HBsAb did not differ at baseline between High and Low Responder cohorts, their humoral trajectories diverged over the course of vaccination. This divergence was most prominent in the HBVnaive-Alum group, where Low Responder titers did not rise post second or third vaccinations (P=3.1e-4, linear mixed effects (LME) interaction; **Fig. 1D**), with a similar phenomenon observed in HBVimmune (P=2.5e-9, LME interaction; **Fig. 1D**). The responder category was not associated with differences in age, sex, body mass index, or smoking status (all P>0.05, multivariate logistic regression; **Fig. S1C**). Overall, these results establish a clinical cohort in which vaccine regimen and pre-existing HBV immunity shape the magnitude of humoral responses, but antibody titer stratification into High and Low Responders revealed pharmacodynamic differences in response kinetics that were not explained by demographic or clinical variables alone.

### Ipsilateral lymph node enlargement following vaccination

To uncover immune mechanisms associated with humoral responses, we directly sampled the vaccine-draining axillary lymph node following vaccination. Of the 86 individuals offered optional CNBx, 72 (83.7%) consented to ultrasound screening, of whom 10 (13.9%) had lymph nodes amenable to ipsilateral ultrasound-guided biopsy at a mean of 15.6 (SD 3.6) days after the final vaccination visit. Ultrasound measurements of lymph nodes were a mean area of 0.79 cm^2^, long axis length of 1.43 cm and cortex thickness of 0.29 cm (**Table S2, Fig. 1E and S1D**), measurements similar to those reported following SARS-CoV2 vaccination (*37*, *38*). We considered whether participants with lymphadenopathy differed from participants without. However, CNBx participants did not have uniformly strong humoral responses and their distribution of antibody responses was similar to that of the overall cohorts (**Fig. 1C**). The CNBx subgroup also reflected the overall study age distribution (N.S., Wilcoxon; **Fig. S1E**). Although addition of CpG adjuvant improves antibody responses in prior studies (*39*, *40*), lymph nodes were not consistently enlarged in HBVnaive-CpG recipients, as the rates of CNBx were 11% (n=1) for HBVnaive-CpG vs 8% (n=3) for HBVnaive-Alum, and 22% (n=6) for HBVimmune (**Fig. S1F**). We also considered participants’ recent vaccine history, as GC reactions following vaccination can be prolonged (*41*), but only one participant had had any recorded vaccination in the 6 months prior to study entry, and laterality could not be established. Thus, the CNBx subset appeared representative of the larger cohort, with respect to the clinical parameters recorded in the study.

We next evaluated the CNBx specimens histologically. One core per individual was formalin-fixed and paraffin-embedded (FFPE) for histology while remaining cores were mechanically dissociated and cryopreserved. To identify GCs, we first optimized immunofluorescence (IF) staining for Ki67, IgD, and PD-1 in unrelated, deidentified tonsil tissue. Hematoxylin and eosin (H&E) staining revealed that six of ten FFPE sections were consistent with lymphoid tissue, and subsequent IF detected GCs in four of these samples. These GCs displayed the expected polarization of Ki67 and PD-1 though were smaller than tonsil GCs (P<0.05, adjusted Wilcoxon; **Fig. 1F-G**). The absence of GCs in the remaining six samples likely reflects sampling error, as only a single core per participant was preserved for histology and GC cells were identified by single cell analysis (**Fig. S2A**). These data confirmed the presence of GCs in ipsilateral lymph nodes following HBV vaccination.

### Germinal center B cell activity and transcriptional programs distinguish robust humoral responses

To investigate the cellular and clonal dynamics underlying divergent humoral responses in our High and Low Responder groups (**Fig. 1C-D)**, we performed single-cell RNA sequencing on lymph node cells from all 10 CNBx participants. In the B cell compartment, we recovered 22,037 B cells in total (**Fig. 2A**). Unsupervised clustering resolved six distinct B cell populations according to established marker genes (*27*, *42*, *43*): dark zone (DZ) and light zone (LZ) GC-B cells, antibody secreting cells (ASC), atypical B cells, and two mixed phenotype clusters (Mixed_B1 and Mixed_B2) (**Table S3; Fig. 2A-B and S2A-D**). BCR repertoire profiling revealed higher IGHV mutation frequency in ASC and GC-B cell clusters when compared to mixed B cell subsets (Wilcoxon, P<0.05; **Fig. S2B**). Lymph node B cell composition varied by participant (**Fig. S2C**) but showed no substantial differences between immune and naive cohorts (**Fig. S2E**). Next we evaluated whether the frequency of the GC-B cell subsets was different between these responder groups. Indeed, miloR (*44*) differential abundance analysis demonstrated LZ GC-B cell enrichment in High Responders relative to Low Responders in 85% (302/355) of iterations with detectable differences (**Fig. 2C**).

**Figure 2.**
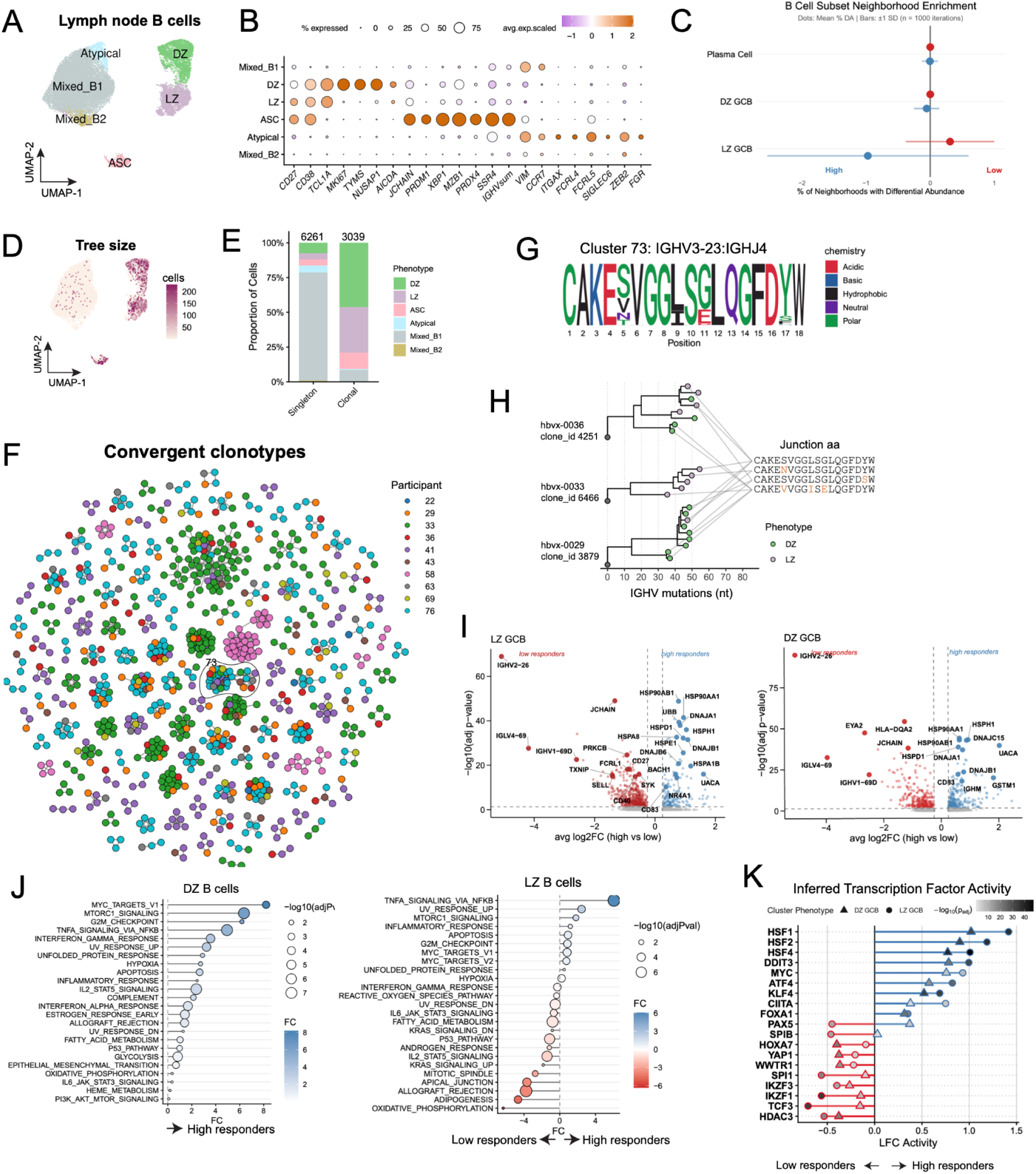
Germinal center B cell activity and transcriptional programs distinguish robust humoral responses. **A.** UMAP of lymph node B cells from all CNBx participants, resolving six populations by unsupervised clustering: Atypical (light blue), Mixed_B1 (grey), Mixed_B2 (golden brown), dark zone (DZ) GC-B cells (green), light zone (LZ) GC-B cells (light purple), and antibody-secreting cells (ASCs, pink). **B.** Dot plot of key lineage and functional marker genes used to define and annotate each B cell subset. Dot size reflects percent of cells expressing the gene and color reflects scaled average expression. **C.** Differential abundance (% DA) of cellular neighborhoods for ASCs, DZ and LZ B cells stratified by High (blue) and Low (red) Responders. Dot indicates mean percent differential abundance and horizontal bars represent SD across 1000 iterations. **D.** Distribution of clonal lineage tree size projected onto the B cell UMAP, highlighting where expanded clones are concentrated. **E.** Proportion of B cell phenotypes among singleton (unexpanded) versus clonal (expanded) BCR repertoires. Numbers above bars denote total B cells in each category. **F.** Network of heavy chain convergent clonotypes shared across participants, where each cluster of nodes shares IGHV and IGHJ gene usage and ≥85% CDR3 amino acid sequence identity. Nodes are colored by participant ID. Outlined cluster 73 is detailed further in G and H. **G.** Sequence logo of the CDR3 junction amino acid sequence consensus for cluster 73, a convergent clonotype cluster outlined in F. Cluster 73 cells share IGHV3-23 and IGHJ4 gene usage. Residues colored by side chain chemistry. **H.** Clonal lineage trees for representative expanded clones within cluster 73 (from F), drawn from three separate participants (hbvx-0036, hbvx-0033, hbvx-0029). Nodes colored by B cell phenotype (DZ or LZ), and gray connecting lines link tree tips to their shared CDR3 junction sequence. **I.** Volcano plots of differentially expressed genes between High and Low Responders within the LZ and DZ GC-B cell subsets. **J.** Single-cell pathway analysis (SCPA) of MSigDB Hallmark gene sets comparing High and Low Responders within the DZ and LZ GC-B cell compartments. Dot size reflects statistical significance (−log10 adjusted P) and color reflects fold change. **K.** Inferred transcription factor activity (decoupleR) for DZ (triangles) and LZ (circles) GC-B cells, expressed as log-fold change (LFC) between High and Low Responders. Point shading reflects −log10 adjusted P.

To resolve clonal relationships in GC populations, we mapped lineage and clonotype dynamics across subsets and participants (**Fig. 2D**). Expanded clones represented a minority of the total captured B cell pool (**Fig. S2F**) but were enriched within the GC compartment (**Fig. 2D-E**), whereas nonexpanded repertoire mapped almost exclusively to non-GC B cells (**Fig. 2E and S2F**). We next investigated whether vaccination elicited convergent repertoires across individuals. Convergence was defined by matching IGHV/IGHJ usage, junction length, and amino acid sequence similarity of ≥85% (**Fig. 2F**), yielding 129 clusters, of which 93 (72%) were present in at least two participants and 47 (36%) were shared by three or more (**Fig. 2F and S2G**). For example, a consensus CDR3 sequence for one of these clonotypes (CAKESVGGLSGLQGFDYW) was identified across GC-B cells in eight different individuals (**Fig. 2G-H**). The convergent repertoire consisted of predominantly GC-B cells and ASCs (**Fig. S2G-H**). Moreover, participant lineage trees containing a clonotype shared with at least one other participant, denoted as shared repertoire, demonstrated greater mutational burden (Wilcoxon, P = 1.2e-11; **Fig. S2I**), longer HCDR3 sequences (Wilcoxon, P=2.1e-4; **Fig. S2J**), and were enriched for *IGHV3-23*, *IGHV1-69*, and *IGHV4-34* in shared relative to non-shared repertoire (**Fig. S2K**), of which *IGHV3-23* has been previously reported in HBV vaccine responses (*45*). Within the convergent pool we identified public clonotypes across 8 of 10 individuals comprising 1726 B cells which were predominantly DZ and LZ GC-B cells or ASCs (**Fig. S2L**). Within responder group (High-High, Low-Low) number of public clonotypes did not differ significantly from across-group (Low-High, High-Low) sharing in either responder groups (High P= 0.1; Low P=0.12, Wilcoxon signed-rank test; **Fig. S2M**), suggesting that divergent antibody response was not due to a failure to recruit relevant B cells.

To assess whether these shared clonotypes aligned with known specificities, we cross-referenced HBV CNBx repertoires with prior studies (*45–47*). We identified 5 clonotypes in 3 CNBx participants that converged with clonotypes found in published HBV vaccine-specific responses (*45–47*)(**Table S4**). Moreover, the immune epitope database (IEDB) matched only one clonotype across all participants, in this case for SARS-CoV-2 spike glycoprotein binding (*48*), indicating a low background rate of coincidental matches relative to the clonotypes identified above.

We next sought to identify the cellular and transcriptional elements associated with variable antibody responses, using the High and Low Responder stratification established in **Fig. 1C-D**. We considered whether B cell mutational burden was similar between the two strata. Indeed, DZ and LZ IGHV mutations were higher in High Responders when compared to Low Responders (**Fig. S2N**). To identify transcriptional programs underlying these two strata, we performed differential gene expression analysis of GC B cell subsets (**Fig. 2I, Table S5**). In High Responders, DZ GC-B cells had greater expression of heat shock protein family members (*HSP90AA1, HSPH1, HSP90AB1, HSPD1*) as well as co-chaperones (*DNAJC15, DNAJA1, DNAJB1*), which may support *BCL6* in the DZ compartment (*49*). In the LZ compartment, High Responders similarly upregulated heat shock protein family members (*HSP90AA1, HSPH1, HSPA1B, HSP90AB1, HSPD1, HSPA8, HSPE1*) and DNAJ protein family members (*DNAJA1, DNAJB1, DNAJB6*), while Low Responders were enriched for *SELL, SYK, FCRL1,* and *JCHAIN*, a transcriptional profile inconsistent with committed GC programming (*50*). Here, DZ and LZ gene expression differences delineate a divergence; GC-B cells in High Responders are transcriptionally active, whereas Low Responders, GC-B cells present a more quiescent state. Pathway analyses confirmed inference from individual transcripts. The DZ GC-B cells from High Responders had increased pathway activity for MYC targets, NFκB signaling, mTORC1 signaling, and G2M checkpoint, indicative of more active metabolic engagement and productive GC cell cycling. The LZ compartment exhibited a divergence between High and Low Responders (**Fig. 2J**). High Responder LZ GC-B cells were also differentially enriched for TNF/NFκB signaling, Inflammatory response, mTORC1 signaling, and MYC target gene sets. This profile reflects non-canonical NFκB signaling, downstream of CD40, which promotes MYC upregulation in the LZ GC-B cells (*51*, *52*). However, Low Responders also had enrichment of the P53 Pathway, which may indicate cell cycle arrest, as well as IL2/STAT5 signaling which regulates BCL6 (*53*). Taken together, these data suggest that LZ B cells from High Responders were highly receptive to positive selection signals from Tfh, whereas Low Responder LZ B-cells instead remained in a more metabolically-quiescent or stalled state.

Having identified differential gene and pathway expression, we next sought to determine whether these differences were underpinned by distinct regulatory programs. We inferred transcription factor activity networks using decoupleR (*54*) within LZ and DZ compartments independently and stratified by High and Low Responder groups. High Responders displayed a consistent proteostatic and proliferative signature spanning both compartments, marked by heightened activity of heat shock and unfolded protein response regulons *HSF1*, *HSF2*, and *DDIT3*, alongside *MYC* (**Fig. 2K**). This proteostatic program was pronounced in the LZ, where High Responders additionally exhibited elevated ATF4 and *CIITA* activity, consistent with a coordinated transition toward productive and active responses including antigen presentation and antibody production. In contrast, Low Responders instead exhibited a subdued profile more consistent with early activation. In the DZ, this was characterized by elevated Hippo activity (*YAP1*, *WWTR1*) (*55*, *56*). In the LZ, Low Responders aberrantly sustained B cell identity programs (*PAX5, SPI1, SPIB*, and *IKZF3*). These data support a model in which High Responders execute a coordinated GC program, whereas Low Responders fail to dynamically navigate either regulatory checkpoint.

To corroborate these inferred regulatory landscapes, we utilized single-cell regulatory network inference and clustering (SCENIC) as an orthogonal regulon analysis (*57*, *58*) (**Fig. S2O, Table S6**). Within the LZ compartment, High Responders demonstrated dominant *RELB* and *NFκB2* activity, drivers of the non-canonical NFκB pathway activated downstream of CD40. Stress responsive regulons *ATF4, ATF3, CEBPB,* and *CEBPG* may reflect productive GC-B cell responses, while *IRF1* and *IRF9* activity are aligned with heightened inflammatory and interferon signaling observed at the pathway level. Conversely, Low Responder LZ GC-B cells demonstrated dominant *IRF2, PAX5, STAT2,* and *BCL6* regulon activity. Rather than engaging the costimulatory selection programs observed in High Responders, Low Responder LZ B cells had persistence of basal identity factors (PAX5, BCL6) alongside IRF2-driven suppression of interferon response pathways.

### T follicular helper subsets are spatially segregated

The enrichment and active state of LZ GC-B cells in High Responders suggested that these B cells were undergoing interaction with LZ-positioned Tfh and identified the LZ region and T cell help as most relevant for investigating determinants of differential humoral response. To understand whether the Tfh programs differ between response groups, we first performed unsupervised clustering and UMAP visualization of the CD4+ T cells recovered from CNBx. The cluster analysis revealed distinct populations of naive, memory, regulatory, and Tfh cells (**Fig. 3A and S3A**; **Table S3**). To evaluate for the presence of HBV-specific CD4+ T cells, we mapped HBV-specific CD4+ T cell clonotypes by cross-referencing CDR3 sequences with known HBV epitopes from a study of chronic HBV infection (*59*) in the IEDB database. In total, we identified 44 matching clonotypes across all CD4+ T cells (P=8.1e-96, Fisher’s Exact test) including 4 in the Tfh cluster (P=7.4e-6, Fisher’s Exact test, **Fig. 3B, Fig. S3B**), indicating the presence of antigen-specific lymphocytes in the Tfh cluster (*59*).

**Figure 3.**
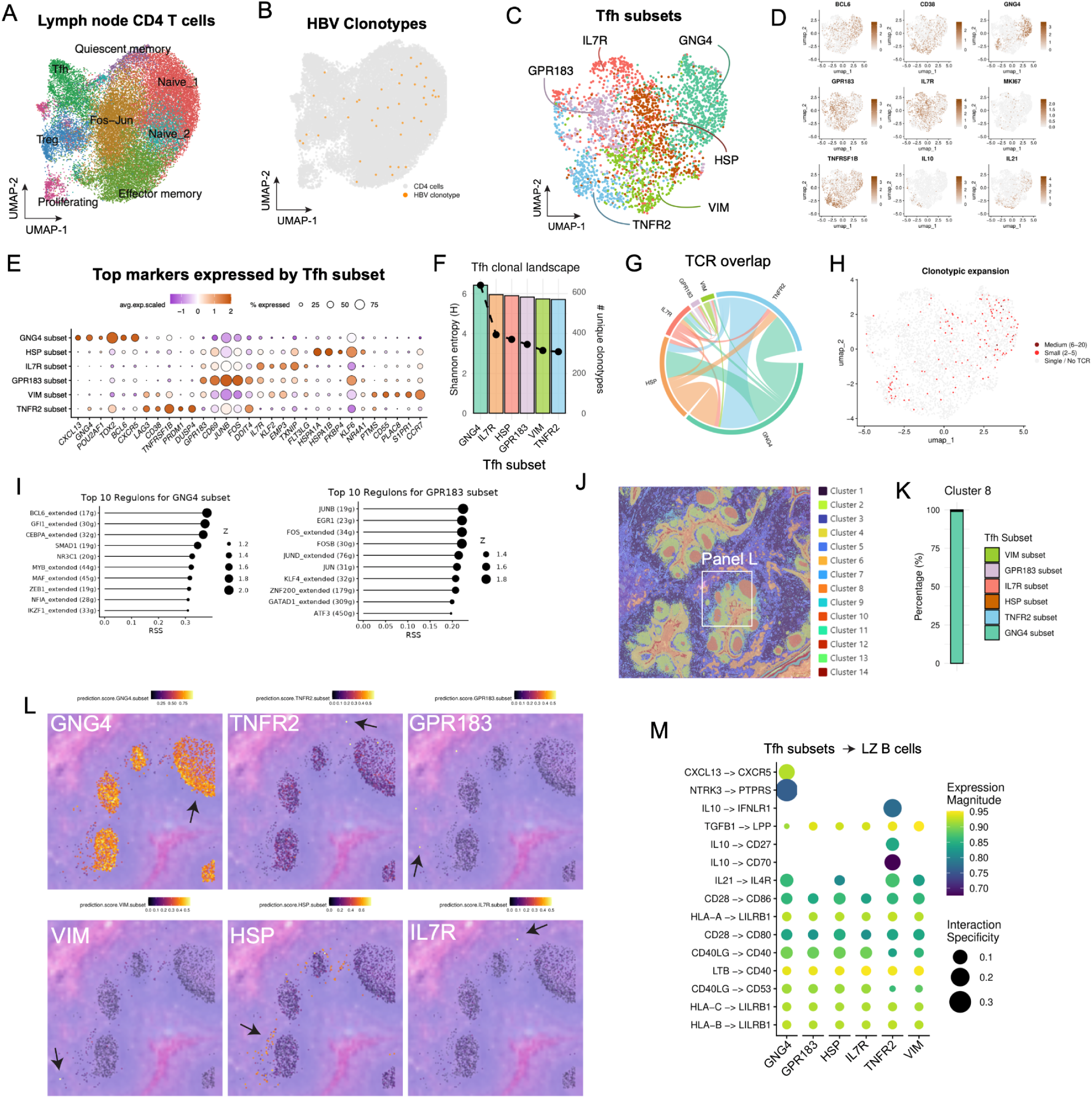
T follicular helper subsets are spatially segregated. **A.** Total lymph node CD4+ T cells, colored by phenotypic clusters: Tfh (green), Treg (blue), Proliferating (magenta), Fos-Jun (mustard), Naive_1 (coral), Naive_2 (teal), Effector memory (olive), Quiescent memory (purple). **B.** Distribution of hepatitis B virus (HBV) specific T cell clonotypes (orange) overlaid on the total CD4 T cell landscape (gray). **C.** Subclustering of the Tfh population identifies six transcriptionally distinct subsets, each named for a uniquely enriched marker gene: IL7R, GNG4, GPR183, TNFR2, VIM, and HSP. **D.** Feature plots showing normalized expression of canonical Tfh and functional marker genes across total Tfh population. **E.** Dot plot of top marker genes defining each Tfh subset. Circle size denotes the percentage of cells expressing the gene and color scale denotes relative average expression. **F.** Clonal diversity of the six Tfh subsets, indicating Shannon entropy values (bars, left y-axis) and total unique clonotype count (line, right y-axis). **G.** Chord diagram of TCR clonotype overlap and clonal sharing dynamics between Tfh subsets. **H.** Clonotypic expansion within the Tfh population, categorized into Medium (6–20 cells), Small (2–5 cells), and Single/No TCR groups. **I.** Regulon specificity score (RSS) and Z-score rankings of the top 10 transcription factor regulons defining the GNG4 subset (left) and GPR183 subset (right), inferred by SCENIC. **J.** Clustering of a reference human tonsil section spatial dataset (Visium HD) with 14 microanatomical clusters. White box delineates enlarged representative region (Panel L). **K.** Composition of Tfh subset identity within microanatomical cluster 8 (predominantly GC). **L.** Spatial transcriptomic deconvolution and prediction scores for each Tfh subset (GNG4, TNFR2, GPR183, VIM, HSP, and IL7R) projected onto tonsil tissue section. Black arrows indicate regions of specific spatial enrichment. **M.** Predicted ligand-receptor interactions between Tfh subsets (source) and LZ B cells (target), inferred by LIANA. Color intensity reflects the expression magnitude of the interaction pairs, and circle size denotes NATMI-derived interaction specificity.

We next focused on the Tfh compartment, which we identified by unsupervised clustering of CD4+ T cells and expression of canonical markers including *CXCR5*, *BCL6*, and *CXCL13*. We performed further subclustering of these 2,995 Tfh cells, as it has become increasingly recognized that Tfh subsets may have distinct properties (*15*, *60*, *61*). Subclustering of the Tfh cluster revealed six distinct subsets (**Fig. 3C-D and S3C**). These subclusters were named according to gene expression patterns (**Fig. 3D-E, S3D**). The subset associated with the classical germinal center Tfh state was designated the GNG4 subset, as described in a recent study (*15*), and was characterized by co-expression of *CXCL13*, *POU2AF1*, *TOX2*, *BCL6*, and *CXCR5*. Another subset, termed the GPR183 subset, was marked by the expression of *GPR183*, *CD69*, *JUNB*, and *FOS*. We also identified a TNFR2 subset expressing early activation genes including *LAG3*, *CD38*, *TNFRSF1B*, *IL10*, and *PRDM1*; an IL7R subset indicative of a quiescent or memory-like state (*IL7R*, *KLF2*, *TXNIP*); a VIM subset characterized by structural and migratory markers (*PTMS*, *S1PR1*, *CCR7*); and an HSP subset enriched for stress response genes (*HSPA1A*, *HSPA1B*). All subsets had relatively similar Shannon’s entropy (**Fig. 3F and S3E**). TCR overlap analysis showed substantial sharing of clonotypes between the GNG4, TNFR2, and GPR183 states (**Fig. 3G**), suggesting potential transitions between these functional states within the lymph node. Furthermore, clonotypic expansion, which was defined as observation of two or more clones of the same clonotype, was most visible in the GNG4 and TNFR2 subsets (**Fig. 3H**). Finally, to infer potential developmental relationships between these Tfh subsets, we performed unsupervised trajectory analysis using Slingshot (*62*). Pseudotime ordering, without establishing a root node, arranged the cells along a continuous differentiation trajectory, indicating the GNG4 subset was likely more transcriptionally polarized than other subsets (**Fig. S3F-G**). Other subsets such as GPR183 and IL7R had intermediate pseudotime values, whereas the GNG4 subset had the greatest pseudotime values which may reflect progression towards the GC-Tfh state.

To delineate the regulatory networks orchestrating these diverse transcriptional profiles, we performed SCENIC (*57*, *58*) (**Table S6**). By evaluating transcription factor binding motifs alongside co-expression modules, we identified unique regulons driving the specialization of each Tfh state (**Fig. 3I and S3H-I**). The GNG4 subset was governed by an expected regulatory program driven by master transcription factors including *POU2AF1* and *BCL6*, in accordance with prior studies characterizing broader Tfh identity (*63*). In contrast, the GPR183 subset was instead dominated by a marked activation of the AP-1 complex network, exhibiting high specific activity for *JUN*, *JUND*, *FOS*, *FOSB*, *EGR1,* and *JUNB*, which suggests the GPR183 subset represents a highly activated, early-primed state. In contrast, the TNFR2 subset was associated with transcription factors linked with terminal differentiation and activation-induced dampening, heavily utilizing *PRDM1* (Blimp-1), *CREB3L2*, and *BHLHE40* regulons which are known to regulate Tfh programs (*64*, *65*). Conversely, the *IL7R* subset was actively maintained by canonical T cell memory and stemness regulators, such as *KLF2* which also have been associated with restraint of the Tfh program (*66*). Finally, the stress-associated *HSP* subset was transcriptionally isolated by its reliance on heat-shock factors, specifically *HSF1*, *HSF2*, and *CEBPB*. Together, these distinct regulon architectures detail the gene regulatory networks involved in GC Tfh.

We next sought to identify the physical location of Tfh transcriptomic subsets within the intact lymphoid architecture to determine which would be most likely to be involved in the germinal center reaction. We mapped the transcriptional signatures of our Tfh subsets onto a reference dataset of a human tonsil with spatial clustering generated using Visium HD (**Fig. 3J-L and S3J-L**). One of these spatial clusters, Cluster 8, was identified as the mature germinal center core based on dense *BCL6* expression. Quantitative composition analysis of the Cluster 8 spatial bins revealed that the *GNG4* signature constituted nearly all of the Tfh signal within the GC niche. Conversely, the TNFR2 and GPR183 subsets mapped predominantly to the extrafollicular space and the T-B border regions, thereby corroborating their potential roles as activated and pre-Tfh transitional states in the setting of immature GCs.

Based on the evident proximity of the GNG4+Tfh with LZ-B cells, we evaluated receptor-ligand interactions of these two subsets using LIANA (*67*). The GNG4+Tfh subset was the only Tfh subset that demonstrated *CXCL13-CXCR5* interaction (**Fig. 3M**), consistent with prior literature (*68*), as well as interactions such as *NTRK3-PTPRS* known to be regulated by STAT3 (*69*). Together, these data support the notion that the GNG4+Tfh subset is the primary subset interacting with LZ-B cells in the germinal center.

### GNG4+ Tfh cell activity is a hallmark of humoral responses to HBV vaccination

Having mapped the CD4+ T cell landscape and defined discrete Tfh functional states, we next sought to determine how these subsets correlated with vaccination outcomes. We first evaluated compositional shifts in the Tfh compartment relative to vaccine response. Although there were no substantial differences in the proportion of Tfh subsets within High and Low Response (**Fig. S4A**), Cohen’s *d* effect size calculation identified the GNG4+Tfh subset as having the strongest association with High Responders (**Fig. 4A, S4A**). Indeed, the odds that the GNG4+Tfh subset was more likely to be associated with High Responders was over 10.1 times higher than for Low Responders, followed by the TNFR2 subset at 5.2 times higher. We also performed miloR differential abundance testing to evaluate enrichment of the different clusters, which demonstrated that the GNG4+Tfh subset was most reliably associated with High Responders, followed by the GPR183 subset (**Fig. 4B-C and S4B**). Notably, several subsets were associated with Low Responders, such as the HSP and IL7R subsets, which may reflect a failure to fully engage mature, follicle-resident helper cells. We thus focused on the GNG4+Tfh subset given its spatial position in the GC and its effect size.

**Figure 4.**
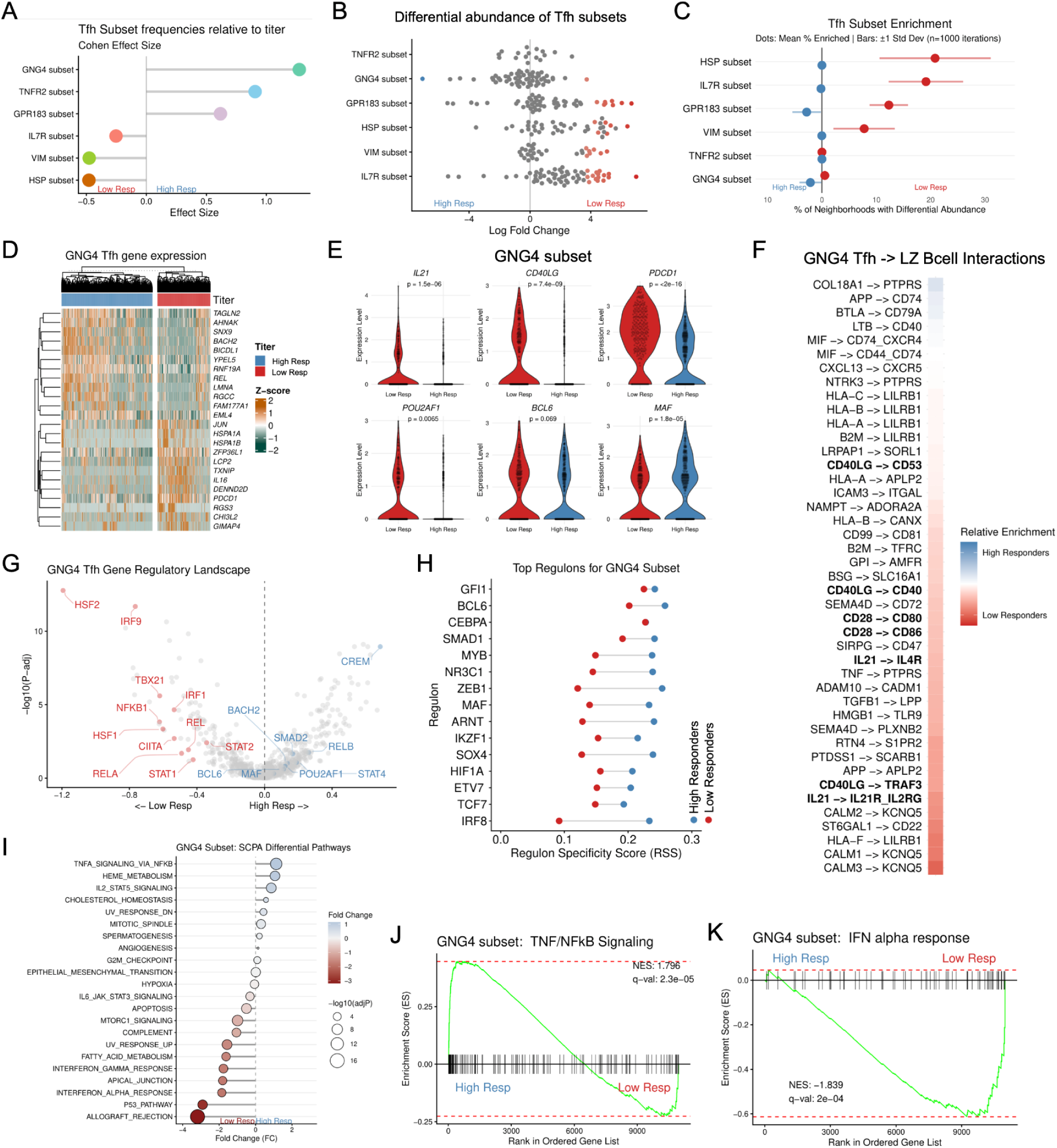
Transcriptional programs in GNG4+Tfh cells associate with humoral responses to vaccination. **A.** Cohen’s effect size for frequencies of lymph node Tfh subset (GNG4, TNFR2, GPR183, IL7R, VIM, HSP) between High and Low Responders. **B.** Log fold-change distribution of Tfh subset abundance across single-cell neighborhoods, comparing High versus Low Responders. Colored points denote neighborhoods with significant differential abundance. **C.** Percentage of neighborhoods per subset showing significant differential abundance by miloR. Dots represent mean percentage enriched, and bars indicate ±1 standard deviation across 1,000 iterations. **D.** Unsupervised hierarchical clustering of differentially expressed genes within the GNG4+Tfh subset, stratified by High and Low response groups. **E.** Single-cell expression for selected genes within the GNG4+Tfh subset, comparing Low and High Responders. P value by Wilcoxon test. **F.** Reciprocal ligand-receptor interactions from the GNG4+Tfh subset (source) to LZ GC-B cells (target), inferred by LIANA. Relative enrichment reflects the difference in aggregate_rank scores between High and Low Responders. **G.** Volcano plot of differential gene regulatory network activity within the GNG4+Tfh subset (decoupleR), comparing Low and High Responders. **H.** Regulon Specificity Scores (RSS) for the top 10-ranked transcription factor networks governing the GNG4+Tfh subset across each of the High and Low response groups. **I.** Single-cell pathway analysis (SCPA) for differential activation of Hallmark pathways within the GNG4-subset between response groups. **J-K**. Gene set enrichment analysis profiles for the (**J**) TNF/NF-κB signaling and (**K**) IFN-α response pathways.

Although the abundance of GNG4+Tfh cells was associated with high HBsAb titers, we also considered that qualitative, cell-intrinsic functional deficits might also contribute to suboptimal vaccine responses. To address this, we performed differential gene expression and pathway enrichment analyses restricted to the GNG4+Tfh compartment, comparing cells derived from High versus Low Responders (**Fig. 4D and S4C; Table S5**). The top genes associated with GNG4+Tfh from High Responders included *BACH2* and *REL*, whereas the top genes in Low Responders included *PDCD1*, *JUN*, and *TXNIP*. We then evaluated per-cell gene expression for genes characteristically associated with Tfh function, which demonstrated greater expression of *IL21*, *CD40LG*, and *POU2AF1* in GNG4+Tfh from Low Responders (**Fig. 4E**). We next modeled intercellular ligand-receptor interactions to determine if the pattern of differential expression of GNG4+Tfh was associated with altered LZ GC-B cell interactions (**Fig. 4F and S4D-E; Table S8**), recognizing that the primary function of GC Tfh cells is to provide cognate help to B cells. Differential interaction network modeling using LIANA (*70*) to evaluate GNG4+Tfh and LZ B cells demonstrated enrichment for interactions including *CD40LG-CD40, IL21-IL21R*, and *CD28-CD80* in the Low Responders (**Fig. 4F**). Thus, the functional deficit in Low Responders does not stem from a failure to express critical Tfh ligands or form receptor-ligand contacts, nor was it associated with loss of core Tfh transcriptional programs.

To further interrogate the transcriptional gene regulatory network of GNG4+Tfh, we inferred the single-cell regulatory networks using gene regulatory network analysis via decoupleR (*54*). The regulatory landscape of High Responders was anchored by classical helper and maintenance transcription factors, including heightened activity of the *BCL6, SMAD2, CREM*, *STAT4*, and *MAF* regulons despite reduced gene expression of a select few (e.g. *POU2AF1*) (**Fig. 4G; Table S7**). In contrast, the regulatory architecture of GNG4+Tfh cells in Low Responders was marked by stress-induced and inflammatory transcription factors including *STAT1*, *RELA*, *NFκB1*, *IRF9*, and *CIITA*. The heightened activity of these Type I and II interferon-responsive factors indicates an inflammatory rewiring that competes directly with canonical Tfh gene regulatory networks. We also performed SCENIC regulon analysis (*57*, *58*) (**Fig. 4H and S4F**), which also demonstrated greater specific regulon activity for *BCL6* and *MAF* in GNG4+Tfh from High Responders than from Low Responders. Furthermore, gene regulatory networks for AP-1 complex members such as JUN and FOS were notably enriched in the Tfh cells of Low Responders (**Fig. S4F**). These data implicate transcriptional network “noise” in the GNG4+Tfh from Low Responders as associated with poor antibody responses, in spite of expression of *IL21*, *CD40LG*, *BCL6*, and other elements of the canonical Tfh transcriptional program.

To further interrogate the transcriptional networks that distinguished the Responder cohorts, we performed pathway analysis using the MSigDB Hallmark gene sets (*71*). Single-cell pathway analysis (SCPA) revealed a transcriptional rewiring of GNG4+Tfh cells in Low Responders (**Fig. 4I**). In contrast, GNG4+Tfh cells from the Low Responders cohort displayed potent inflammatory and stress-response signatures, including “Interferon Alpha Response” and “Interferon Gamma Response” gene sets. In contrast, TNF/NFκB signaling was associated with High Responders’ GNG4+Tfh, underscoring a central role for NFκB signaling in Tfh help (*22*). GSEA demonstrated positive normalized enrichment score (NES) for “TNFa Signaling via NFκB” (NES = 1.796, q-val = 2.3e-05) and negative “Interferon Alpha Response” (NES = –1.84, q-val = 2e-4), supporting pathway results above (**Fig. 4J-K**). Similarly, the transitional GPR183 subset in High Responders exhibited parallel dysregulation, with enrichment for hypoxia, apoptosis, and TGF-beta signaling pathways (**Fig. S4G**). Additionally, the TNFR2 subset demonstrated similar indicators of dysregulation with enrichment of hypoxia and TNF/NFκB signaling pathways (**Fig. S4H**). Taken together, our results demonstrate that poor humoral responses to HBV vaccination are associated with an inability to quantitatively expand the GNG4+Tfh pool and a qualitative, cell-intrinsic molecular dysfunction characterized by interferon-associated hyperactivation that impaired effective T-B intercellular communication within the GC.

## DISCUSSION

In this study, we combined longitudinal clinical profiling of HBV vaccine responses with ultrasound-guided lymph node biopsy to directly interrogate the GC reaction underlying variable human vaccine responsiveness. Our study evaluated cellular observation 10-21 days after vaccination and linked this to end-of-study antibody titers drawn later. By pairing single-cell BCR and TCR repertoire analysis with transcriptomic and regulon profiling of B and T cell compartments from the same draining lymph nodes, we find that divergent antibody outcomes are not easily explained by a failure of GC formation, convergent antigen recognition, or absence of T cell help. Rather, Low Responders mount a qualitatively and quantitatively distinct Tfh program: a contracted, transcriptionally rewired GNG4+ GC-Tfh population that retains some canonical helper features yet fails to sustain the gene-regulatory architecture required for productive LZ licensing. This suggests specific, molecularly-defined defects are at the center of suboptimal human vaccine responses.

A notable feature of the post-vaccination lymph node was the degree of BCR convergence we observed, both across unrelated participants in our cohort and with previously published HBV-specific sequences. Public or semi-public clonotypes have now been described across a range of human antibody responses, including dengue (*72*), SARS-CoV-2 (*73*), and HIV (*74*). Moreover, convergent BCR evolution has been described in influenza (*75*) and in other vaccination settings as well (*76*), and may reflect the constrained set of BCR that can recognize an immunodominant epitope. We found several of our lymph node clonotypes mapped to published HBV-specific BCR sequences, and, other than a single clonotype which mapped to a SARS-CoV2 glycoprotein epitope, we did not identify any other overlap in the IEDB. Additionally, we did not identify recent vaccinations that might easily explain the substantial pool of public clonotypes and convergent clonotypes across individuals in our study. Moreover, we found that convergence was concentrated in GC-B cell and ASC states and was accompanied by high rates of *IGHV* mutation, which implies an antigen– and selection-driven phenomenon rather than an artifact of naive repertoire sharing. Convergent clonotypic responses did not track with responder status in our cohort, as the recruitment of relevant B cells into the GC occurred even in individuals who ultimately mounted weak antibody titers. This argues that the principal bottleneck limiting humoral output in Low Responders lies downstream of antigen engagement and clonal recruitment rather than at the level of BCR repertoire availability. When considering vaccine design, strategies aimed purely at broadening initial antigen recognition such as via recruitment of diverse founder clones may be insufficient alone without also addressing the cellular processes that sustain GC selection.

Transcriptomic profiling also revealed that High and Low Responders diverged in GC-B cell state. High Responder DZ and LZ B cells were enriched for heat-shock/co-chaperone programs supportive of BCL6 stability, together with MYC and mTORC1 activity consistent with active cyclic re-entry between LZ and DZ compartments. Low Responder GC B cells, by contrast, showed features more consistent with premature extrafollicular commitment or B cell quiescence, including *JCHAIN*, *SELL*, *SYK*, and *FCRL1* expression, a signature associated with early activation or non-GC fates rather than sustained affinity maturation. This divergence was mirrored at the regulon level, where High Responder LZ B cells were governed by *RELB/NFκB2* that can occur downstream of CD40 signaling, whereas Low Responder LZ B cells retained a core identity program including *BCL6* but without the active-selection signatures. These B cell-intrinsic findings suggest that Low Responders adopt some features of canonical GC response but are unable to efficiently progress through iterative cycling in the GC, ultimately raising the question of the contribution of Tfh to this process. Indeed, we found that only a subcluster of Tfh, defined by expression of *GNG4,* is localized to the GC and may represent a candidate cellular biomarker in the lymphoid tissue in humans, consistent with a recent report (*15*). This subset was both quantitatively depleted and qualitatively altered in Low Responders, who showed persistent AP-1 activity (*JUN*/*FOS*) and a regulon signature (*STAT1, RELA, NFκB1, IRF9, CIITA*) more typical of an inflammatory state than a helper program. Furthermore, our regulon analysis shows greater BCL6 and cAMP-responsive element modulator alpha (*CREM*) in the High Responders which are associated with productive GC reactions (*77*, *78*). Paradoxically, GNG4+Tfh from Low Responders had greater expression of *IL21*, *CD40LG*, and *PDCD1*, which suggests these features were unable to overcome the dysfunction leading to poor humoral responses. Rather than indicating enhanced helper capacity, this pattern, which co-occurred with AP-1 activity, suggests a compensatory state. These findings extend the understanding that Tfh cells are not a monolithic population but comprise multiple transcriptionally and functionally distinct maturation states, and that inflammatory microenvironments can skew Tfh differentiation toward non-productive states even when canonical markers are retained.

Our study has several limitations that should inform the design of future studies. First, only a minority of participants had lymph node enlargement amenable to biopsy, and although we did not detect systematic associations between the likelihood of enlargement and available demographic or clinical metadata, we cannot fully exclude sampling bias in the CNBx subset, nor can we rule out that individuals lacking accessible lymphadenopathy differ systematically from those we studied. Second, our cross-sectional biopsy design captures a single time point (10-21 days post-vaccination) and therefore cannot distinguish whether the Tfh alterations we observe reflect delayed GC initiation, premature GC contraction, or a stable dysfunctional state sustained throughout the response. Longitudinal sampling, for example by serial fine-needle aspiration, would help resolve the kinetics of GC and Tfh dysfunction and could reveal whether early inflammatory Tfh states are reversible. Third, our sample size, and in particular the small number of CpG-adjuvanted participants undergoing biopsy, limited our ability to formally test whether adjuvant choice modifies the cellular programs described here, despite the established clinical benefit of CpG adjuvants for HBV vaccine responses. Larger cohorts will be needed to determine adjuvant mechanisms and whether adjuvants can in part overcome Tfh dysfunction. Finally, our data establish a strong association between GNG4+Tfh and antibody outcome, but establishing causality will require additional functional and interventional studies.

Taken together, our data support a model in which productive humoral immunity requires coordinated engagement across both arms of the GC: LZ selection driven by non-canonical NFκB signaling in B cells paired with an intact, spatially-restricted GNG4+Tfh population capable of delivering critical help. Although derived from HBV vaccination, the pathways implicated here, including non-canonical NFκB/CD40 signaling, AP-1-driven inflammatory skewing, and interferon hyperactivation in Tfh, are not vaccine-specific, and parallel defects may underlie poor responses in other chronically inflamed populations, such as in the settings of aging and obesity. Whether Tfh dysfunction reflects a stable, individual-intrinsic trait or a state-dependent response to a given inflammatory context remains an open question that will be essential for designing future rational vaccination strategies.

## Acknowledgements

We are grateful to Drs. Bertram Bengsch, Amy Baxter, Erietta Stelekati, and Alexander Huang for insightful comments and discussion. We thank the NYU Vaccine Center clinicians, nurses, and laboratory staff for advice, technical assistance, and sample processing help. We thank the NYU Langone Health Center for Biospecimen Research and Development (CBRD), Histology and Immunohistochemistry Laboratory (RRID:SCR_018304), Genome Technology Center (GTC), and NYU Center for Genomics and Systems Biology (CGSB). Finally, we would like to thank all the participants who volunteered to contribute to our studies, without whom this work would not be possible.

## Funding

This work was supported by National Institutes of Health (NIH) grants AI158617, AI148574, and AI082630, as well as through funding from the Hevolution Foundation and the American Federation for Aging Research. The Genome Technology Center at NYU Langone Health is supported in part by NYU Langone Health’s Laura and Isaac Perlmutter Cancer Center Support (grant P30CA016087) from the National Cancer Institute. SBD was supported by the University of Pennsylvania Perelman School of Medicine Medical Scientist Training Program T32 (NIGMS GM007170). This research was supported by the Doris Duke Foundation (2021190) and NIAID (K08AI136660) (LAV) and the Parker Institute for Cancer Immunotherapy and V Foundation for Cancer Research, co-sponsoring a Parker Bridge Fellow Award (DAO).

## Contributions

Concept or study design: HMC, ES, XY, TW, LAV, RSH

Funding acquisition: SBK, CRN, DAO, LAV, RSH

Supervision: DAO, LAV, AN, RSH

Acquisition of data: HMC, XY, RS, CM, MT

Investigation: TW, MT, RSH

Analysis: HMC, ES, XY, RS, CM, SBD

Writing – original draft: HMC, ES, RSH

Writing – review & editing: All authors

## Competing interests

None.

## RESOURCE AVAILABILITY

Inquiries for further information, data or resource availability may be directed to and fulfilled by lead contact and corresponding author, Ramin Herati. All scRNAseq and TCR sequences were deposited in Gene Expression Omnibus (GEO) under accession GSE349037 (reviewer token wxgzyqeyfpydtsd). Bioinformatics scripts were deposited in Zenodo under doi 10.5281/zenodo.21440760.

## MATERIALS and METHODS

### Study design and sampling

This study was conducted as a prospective, open-label, partially-randomized evaluation of immunological outcomes of Hepatitis B virus (HBV) vaccination in adults (NCT04674462). Following approval by the NYU Institutional Review Board (protocol 20-01782), written informed consent was obtained from all participants prior to enrollment and study was carried out in accordance with the principles outlined in the Declaration of Helsinki. We recruited a total of 101 ambulatory participants stratified into two main experimental cohorts: a *de novo* vaccination cohort of HBV-naive adults (n=59) and a recall booster cohort of previously immunized, HBVimmune adults (n=42) based on history and medical chart review. One participant in the HBV-naive cohort was unable to comply with study procedures and was thus excluded from the remainder of the study. In the *de novo* cohort, vaccinations were administered in accordance with manufacturer package insert recommendations. To ensure balanced assignment, we used a permuted block randomization strategy with a block size of 10, yielding a 1:4 allocation ratio of 2 CpG-adjuvanted and 8 alum-adjuvanted vaccine recipients per block. Participants in the HBV-naive cohort received either a two-dose series of CpG-adjuvanted vaccine (Heplisav-B, n=13) administered at baseline and at least 4 weeks later, or a standard three-dose series of alum-adjuvanted vaccine (Engerix-B, n=46) administered at baseline, at least 4 weeks, and at least 24 weeks following enrollment. Participants in the HBVimmune cohort received a single booster dose of the alum-adjuvanted vaccine at study entry. Longitudinal peripheral blood sampling was collected at baseline (w0), 1 week after each vaccination (w1), and 4 weeks following each vaccination (w4). All relevant dates were adjusted according to a randomized (but consistent) offset +/− 5 days.

Serum anti-HBsAg antibody titers were quantified using quantitative ELISA kit (XpressBio cat# WB2896) according to the manufacturer’s protocol. Based on the anti-HBV surface antigen titers (HBsAb) at the end of the study, participants were stratified into High Responders (≥100 mIU/mL HBsAb) and Low Responders (<100 mIU/mL HBsAb).

### Primary sample processing

Lymph node samples were mechanically dissociated and filtered via 70 um filter then cryopreserved in fetal bovine serum (FBS, Fisher) with 10% dimethyl sulfoxide (DMSO). Serum was collected using serum separator tubes per manufacturer instructions (SST, BD Biosciences). All aliquots were stored at −80C until use. Lymph node biopsy samples were thawed in RPMI (ThermoFisher) supplemented with 5 mM MgCl2 and 100 IU/mL DNase (Fisher).

### Axillary lymph node core needle biopsy

To gain a direct window into human germinal center dynamics at the active site of secondary lymphoid response, an optional lymph node sub-study was introduced via a protocol modification several months after study initiation. This optional procedure was offered to the next 86 active participants. Participants opting into this procedure underwent ultrasound screening for enlarged ipsilateral axillary lymph nodes between 10 to 21 days after their final immunization. Of the 86 individuals offered this procedure, 72 (83.7%) consented to screening, from whom 10 individuals (HBVnaive-Alum, n=3; HBV-naive-CpG, n=1; HBVimmune-Alum, n=6) presented with lymph nodes amenable for biopsy. Ultrasound-guided core needle biopsies (CNBx) were successfully performed using an 18-gauge needle, yielding one tissue core that was immediately fixed in formalin and embedded in paraffin for immunofluorescence assessment and up to 3 remaining cores that were mechanically dissociated to yield between 3e^5^ – 1e^6^ viable single cells. Single-cell lymphoid suspensions were cryopreserved as described above. Only grade I adverse events were identified during these procedures.

### Immunofluorescence staining

Deidentified tonsils were obtained from NYU Center for Biospecimen Research & Development (CBRD) to serve as GC size comparators. FFPE tissue sections (5 um thick) were mounted and deparaffinized with HistoChoice clearing agent (Fisher) followed by graded ethanol series. Antigen retrieval was performed under high pressure with Dako Target Retrieval solution (Agilent). Sections were first blocked with 5% animal serum, then stained with primary antibody cocktail, consisting of Ki67 (mouse anti-human, clone B56), PD-1 (rabbit anti-human, clone EPR4877), IgD (rat anti-human, clone W18340A), followed by a secondary stain of anti-mouse Cy5, anti-rat Alexa Fluor 488 and anti-rabbit Alexa Fluor 555. Coverslips were applied with ProLong Gold DAPI mountant and cured for at least 24 hours prior to imaging. Fluorescent images were acquired using a 10x objective on Keyence BZ-X810 fluorescence microscope.

### CITEseq

Samples were blocked with Fc receptor blocking solution (Biolegend) and FcR Blocking solution (Miltenyi), then stained for Totalseq-C human hashtag oligos and a panel of surface protein antibodies (Biolegend). Cells were pooled, loaded on Chromium GEM-X microfluidics chips and run on Chromium X controller (10x Genomics). Surface protein, gene expression, and V(D)J immune receptor libraries were constructed with accompanying kits (10x Genomics) and prepared as instructed by the manufacturer. Libraries were quantified by qPCR (NEB Next Library Quantification, Illumina) and pooled for sequencing on the NovaSeq 6000 at the NYU Center for Genomics and Systems Biology core and the NovaSeq X at the NYU Langone Genome Technology Center core.

### Single cell RNA sequencing data processing

CITEseq paired-end FASTQ reads were processed with *cellranger multi* algorithm and aligned to reference human genome GRCh38 with Cellranger v8.0.1 (10x Genomics). Seurat was utilized for single cell library processing and downstream analysis. Quality control filters for cell exclusion included low and high UMI counts, and high mitochondrial counts per processed batch. HTO demultiplexing of genetically distinct individuals was conducted by CLR normalization (*NormalizeData*()), *HTODemux*(), and Single nucleotide polymorphism (SNPs) identification via Souporcell (*79*). Doublets were identified and removed with scDblFinder (v1.20.2), assuming a doublet rate of approximately 0.4% per 1000 cells. Additionally, cells with *CD3E* and *CD19* coexpression were removed. Integration was conducted between different 10x Genomics chip lanes and batches (including normalization with SCTransform, *SelectIntegrationFeatures*() with 5000 features, and *FindIntegrationAnchors*() encompassing the first 20 dimensions, rpca reduction, and 20 k.anchors), followed by subsequent Harmony (v.1.2.4) integration of subclustered cell populations. Celltype annotations were facilitated via Azimuth (v0.4.6).

The previously annotated lymph node CD4+ T cell compartment was subsetted for targeted analysis. To identify distinct T follicular helper maturation states, Tfh cells, defined by canonical expression of *CXCR5*, *BCL6*, *CXCL13*, and *PDCD1*, were isolated and subjected to unsupervised subclustering. Following scaling and normalization, principal component analysis (PCA) was performed, and the top principal components were used to construct a k-nearest neighbor (KNN) graph using the *FindNeighbors()* function (dims=1:30). Community detection was executed via the *FindClusters()* function (resolution=0.5), and dimensionality reduction was visualized using Uniform Manifold Approximation and Projection (UMAP). Cluster-defining marker genes were identified using the Wilcoxon rank-sum test to compare each subset against all other Tfh cells.

Differential expression analysis of the Tfh subsets was performed using *FindMarkers*() function in Seurat, followed by filtering for genes that had an average log2FC > 0.5, had p_adj <0.05, and was expressed in at least 20% of the cells from either of the comparator groups.

### B cell receptor repertoire analysis

V(D)J CITEseq processed files were annotated via IgBlast (v1.22.0) and further formatted using Change-O (v1.3.4) (*80*). Cellular barcodes with multiple heavy chain sequences were excluded. Following per participant Hamming distance clustering, a clonal threshold of 0.17 was established by the gaussian mixture model. Germline sequences were inferred using *createGermlines()* from package Dowser (v2.3), mapping sequences to IMGT human V(D)J references. Proportion IGHV nucleotide mutations relative to germline were calculated using *observedMutations()* from SHazaM (v1.3.1) (*81*, *82*). Using Dowser, sequences were formatted with *formatClones()* and clonal trees generated with *getTree()* by the phylogenetic maximum likelihood (phangorn) method. Trees were visualized with ggtree (v3.10.1). To analyze global sequence convergence, all sequences were clustered by common heavy chain V and J gene, junction length, and junction similarity by Hamming distance threshold of 0.15 estimated by GMM.

### Pseudotime Trajectory Inference

To model the developmental progression of Tfh cells, unsupervised pseudotime trajectory inference was performed using the Slingshot package (v2.14.0). No root state was specifically designated. Cells were ordered along the inferred developmental continuum, and the pseudotime metric was incorporated into the Seurat object.

### Differential Abundance and Humoral Correlation

For differential abundance analysis, *miloR* v.2.6 was used using k=30, d=30 settings (*44*). Because of the downsampling typically used for miloR analysis, we instead performed consensus analysis. For each of 1000 unique seeds (1:1000), 15% of the cells were sampled for the analysis, then measures of central tendency were calculated across all aggregated runs. The magnitude of compositional shifts was evaluated by calculating the log-fold change and determining Cohen’s *d* effect size for each Tfh subset relative to the antibody titer threshold.

### Module Scoring and Pathway Enrichment Analysis

To evaluate transcriptional pathways, cells were scored against the MSigDB Hallmark gene sets (e.g., “h.all.v2026.1.Hs.symbols.gmt”) to evaluate global pathway activity. First, the function *compare_pathways()* from the Single-Cell Pathway Analysis (SCPA) (v. 1.6.2) was performed using default parameters, and P values were adjusted via false discovery rate (FDR) correction. In addition, gene set enrichment analysis was performed using the fgsea package (v. 1.32.4) on the differentially-expressed genes per subset, as determined by *FindMarkers()*, and gene set variation analysis was conducted via the scGSVA package (v0.0.23) and UCell (v2.10.1).

### CD4 T Cell Clonal Analysis

Single-cell T cell receptor (scTCR-seq) data was integrated with the transcriptomic Seurat object utilizing the scRepertoire R package (v1.11.0). For Fisher’s Exact testing, the total number of possible TCR clonotypes was assumed to be 1e8 (*83*). HBV-specific T cell receptor sequences were obtained from Immune Epitopes Database (IEDB) under Reference 1035279. Clonal diversity within each Tfh subset was quantified by calculating Shannon’s entropy. Clonal expansion was categorized by frequency (e.g., single or medium expanded) and overlaid onto the Tfh UMAP embeddings to evaluate the relationship between transcriptional state and clonal burst size.

### Gene Regulatory Network Analysis

Gene regulatory networks were evaluated using *run_ulm()* from the decoupleR package (v 2.12.0) on the Tfh and GC B cell subsets, then results were compared across antibody titer groups using Wilcoxon test followed by FDR adjustment. For the regulatory network reference, we employed CollecTRI (*84*), a comprehensive meta-resource that integrates multiple databases into a directed regulatory network pruned for high-confidence transcription factor-target interactions. In addition, SCENIC regulatory analysis was independently performed using the SCENIC R package (v1.3.1), and was facilitated by the GENIE3 (1.28.0), RcisTarget (v1.26.0), and AUCell (v1.28.0) packages. *InitializeScenic*() utilized human species-specific motif databases. Genes were excluded based on low detection levels and/or cell presence (minCountsPerGene = 3% of total cells, minSamples = 1% of total cells).

### Cell-Cell Communication

Ligand-receptor interactions were calculated with LIANA (v0.1.14). Rankings were based on aggregate scores from applicable methods (CellPhoneDBv2, NATMI, Connectome, SingleCellSignalR, and iTALK).

Significant interactions were defined with a cut-off of 0.05. Interaction network modeling was targeted specifically to evaluate interactions originating from Tfh sub-populations to LZ and DZ B cells. Aggregate_rank metric was used as the summary statistic as calculated by LIANA, where rank=0 would imply strongest enrichment and rank=1 would imply weakest enrichment. The Difference in Aggregate_rank for High Responders and Low Responders was used to determine relative enrichment.

### Visium HD analysis

Tonsil data was downloaded from a publicly-available dataset from a 10 µm-section of a fresh-frozen tonsil from a 21-year-old male using the Visium Human Transcriptome Probe Set v2.0, then aligned and mapped using Space Ranger v3.1.1 (Visium HD Spatial Gene Expression Library, Human Tonsil (Fresh Frozen), 10x Genomics, published 2024-09-17). The Loupe Browser (v9.0) was used to inspect the data. Spatial transcriptomic output aligned to high-resolution H&E images at a bin resolution of 8µm were used. To ensure precise pixel-to-bin mapping, we utilized the *Read10X_Image()* function in Seurat with the platform-specific scalefactors_json.json file. Data were normalized using SCTransform. Dimensionality reduction was performed via Principal Component Analysis then UMAP. Unsupervised spatial clustering, as previously performed by the vendor, was available as part of the dataset and imported into the Seurat object, yielding 14 distinct microanatomical regions (spatial clusters). To integrate the high-resolution Tfh subsets defined in the scRNA-seq analysis with the spatial data, we utilized the Seurat label transfer workflow. The scRNA-seq object containing the annotated Tfh maturation states served as the reference, while the Visium HD dataset was treated as the query. Transfer anchors were identified between the two datasets using the *FindTransferAnchors()* function, and Tfh subset labels (e.g., GNG4, GPR183, TNFR2, VIM) were projected onto the spatial bins via *TransferData()*.

### Statistics

Nonparametric tests were preferentially used throughout using two-tailed tests at α=0.05, unless otherwise indicated. Where paired analyses were performed, the unpaired data points were excluded. Outlier analysis was not performed and thus outliers were not excluded from the primary analyses. Not significant denoted as N.S. Genes were considered differentially expressed at a false discovery rate (FDR) threshold <= 0.05. Prism 9.0 and R v4.4 were used to perform statistical analyses.

## Notes

https://zenodo.org/uploads/21440761

### Competing Interest Statement

The authors have declared no competing interest.

### Summary of Updates

This version of the manuscript has been revised to update the title, revise the text for clarity, and improve figure formatting.

