## supplemental_figures for "Tfh dysfunction is associated with poor responses to HBV vaccination"

**One Sentence Summary:** Lymph node Tfh dysfunction is associated with poor vaccine responses to hepatitis B virus vaccination.

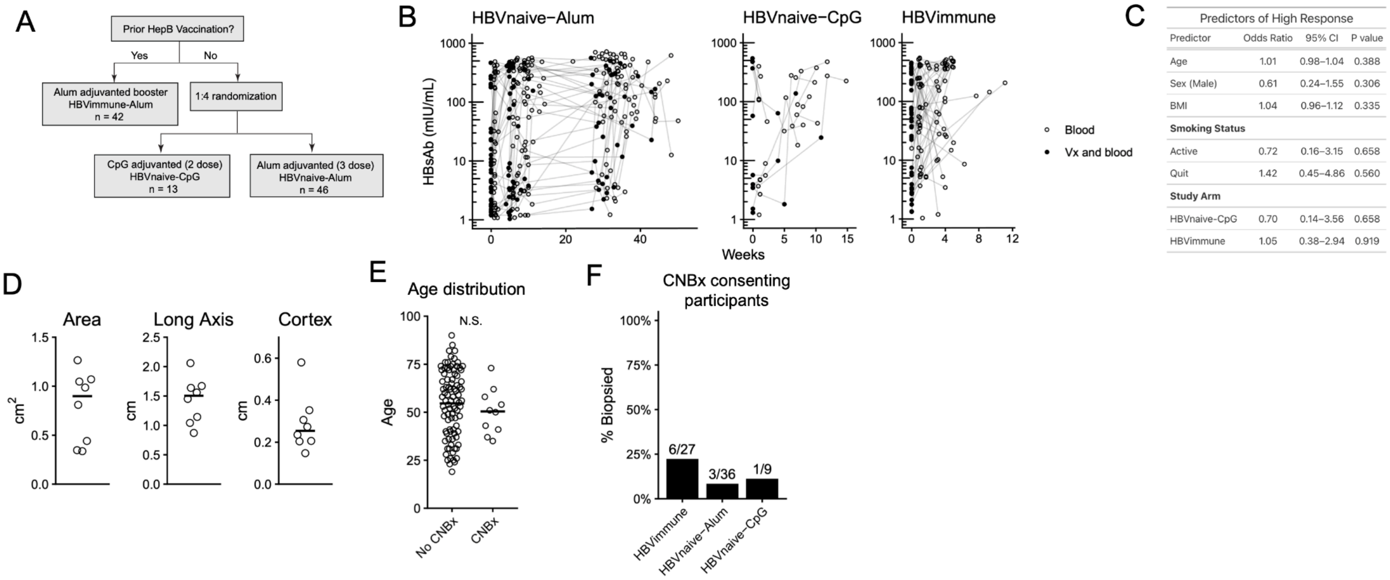


Predictors of High Response

| Predictor | Odds Ratio | 95% CI | P value |
|---|---|---|---|
| Age | 1.01 | 0.98–1.04 | 0.388 |
| Sex (Male) | 0.61 | 0.24–1.55 | 0.306 |
| BMI | 1.04 | 0.96–1.12 | 0.335 |
| **Smoking Status** | | | |
| Active | 0.72 | 0.16–3.15 | 0.658 |
| Quit | 1.42 | 0.45–4.86 | 0.560 |
| **Study Arm** | | | |
| HBVnaive-CpG | 0.70 | 0.14–3.56 | 0.658 |
| HBVimmune | 1.05 | 0.38–2.94 | 0.919 |

**Supplemental Figure 1.**

**A**. Participant randomization and study arm assignment workflow. Individuals with prior vaccination received an alum-adjuvanted booster (HBVimmune, n=42). Vaccine-naive individuals underwent a 1:4 randomization to receive either a 2-dose CpG-adjuvanted vaccine (HBVnaive-CpG, n=13) or a 3-dose alum-adjuvanted vaccine (HBVnaive-Alum, n = 46).

**B**. Longitudinal HBsAb titers by cohort. Individual longitudinal hepatitis B surface antibody (HBsAb) titers (mIU/mL) over time (weeks) across the three study cohorts. White circles indicate blood collection timepoints; black circles indicate concurrent vaccination (Vx) and blood collection events.

**C**. Multivariable logistic regression of clinical and demographic predictors of achieving a high endpoint HBsAb titer. Model includes Age, Sex (ref: F), body mass index (BMI), Smoking Status (ref. Never), and Study arm (ref. HBVnaive-Alum).

**D**. Lymph node anatomical measurements. Quantitative metrics of regional lymph nodes, including cross-sectional area, long axis length, and cortical thickness. Data points represent independent participants, with horizontal bars indicating the mean values.

**E**. Participant ages for all participants and the subset of individuals undergoing CNBx.

**F**. Proportion of participants who successfully underwent CNBx out of the total number screened per study arm.

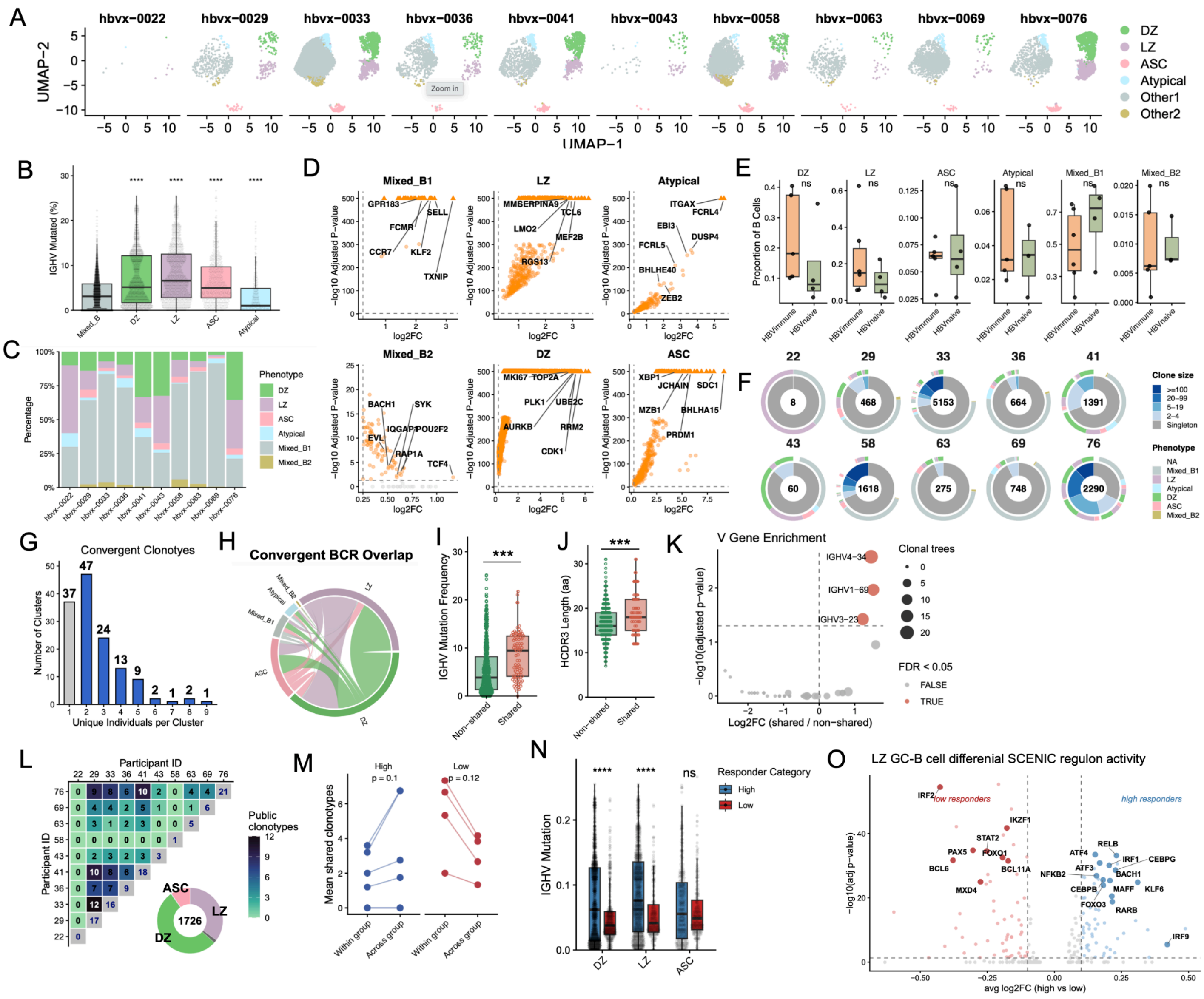


**Supplemental Figure 2.**

**A**. Single-cell UMAP projections of lymph node B cells per individual (light blue=Atypical, grey=Mixed_B1, golden brown=Mixed_B2, green=DZ GC-B cells, light purple= LZ GC-B cells, Pink= ASCs).

**B**. Mutational frequency of each subset compared to aggregate mixed phenotype B cell clusters (Wilcoxon). ****P≤ 0.0001.

**C**. Stacked bar plot of B cell subset composition for each participant.

**D**. Differentially expressed genes between the titled subset and all other B cell subsets. Highlighting top and relevant genes unique to each B cell subset.

**E**. B cell subset proportion by immune and naive categories.

**F**. Distribution of B cell phenotypes (outer ring) and clonal expansion size categories (inner ring). Top number indicates participant and central number denotes total recovered BCR sequences.

**G**. Summary barplot quantifying number of unique participants in each clonotype. Blue bars highlight semi-public clonotypes (each cluster denoted in Fig. 2F) as they are shared between multiple individuals. The number on the bar indicates the number of clonotypes.

**H**. Circos plot of semi-public repertoire as defined in Fig. 2F.

**I**. Mean percentage of mutated IGHV nucleotides between non-shared and shared repertoire. Each point represents the mean of each clonal tree (Wilcoxon, P=1.2e-11).

**J**. HCDR3 amino acid lengths for non-shared and shared clonotypes (Wilcoxon, P=2.1e-4).

**K**. Enrichment of specific IGHV genes (IGHV4-34, IGHV1-69, IGHV3-23) in shared versus non-shared repertoires.

**L**. Pairwise heatmap of public clonotypes between different participant pairs. Donut plot of B cell subset composition of public clonotype repertoire. Center number indicates total number of cells.

**M**. Mean public clonotypes per individual within responder group (High-High or Low-Low) or between responder groups (Low-High) (Wilcoxon signed-rank test).

**N**. Median IGHV mutation by responder groups in the LZ, DZ and ASC compartments.

**O**. Differential transcription factor regulon activity of light zone GC-B cells by SCENIC regulon analysis. N.S., not significant.

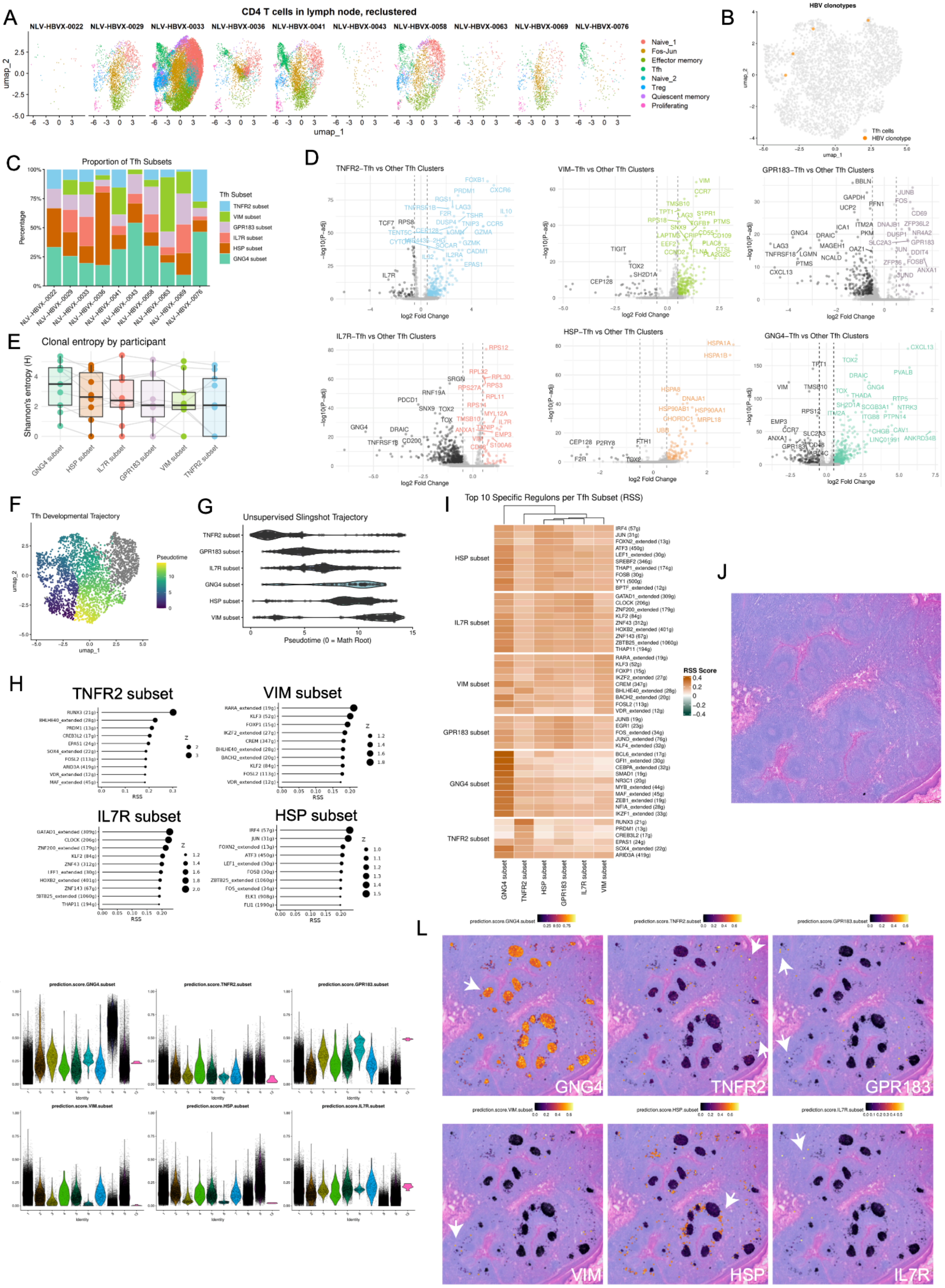

A
CD4 T cells in lymph node, reclustered
NLV-HBVX-0022
NLV-HBVX-0029
NLV-HBVX-0033
NLV-HBVX-0036
NLV-HBVX-0041
NLV-HBVX-0043
NLV-HBVX-0058
NLV-HBVX-0063
NLV-HBVX-0069
NLV-HBVX-0076
umap_1
umap_2
Naive_1
Fos-Jun
Effector memory
Tfh
Naive_2
Treg
Quiescent memory
Proliferating
B
HBV clonotypes
Tfh cells
HBV clonotype
C
Proportion of Tfh Subsets
Percentage
Tfh Subset
TNFR2 subset
VIM subset
GPR183 subset
IL7R subset
HSP subset
GNG4 subset
D
TNFR2–Tfh vs Other Tfh Clusters
VIM–Tfh vs Other Tfh Clusters
GPR183–Tfh vs Other Tfh Clusters
IL7R–Tfh vs Other Tfh Clusters
HSP–Tfh vs Other Tfh Clusters
GNG4–Tfh vs Other Tfh Clusters
log2 Fold Change
–log10(P–adj)
E
Clonal entropy by participant
Shannon's entropy (H)
GNG4 subset
HSP subset
IL7R subset
GPR183 subset
VIM subset
TNFR2 subset
F
Tfh Developmental Trajectory
Pseudotime
G
Unsupervised Slingshot Trajectory
Pseudotime (0 = Math Root)
H
TNFR2 subset
VIM subset
IL7R subset
HSP subset
RSS
I
Top 10 Specific Regulons per Tfh Subset (RSS)
RSS Score
J
K
prediction.score.GNG4.subset
prediction.score.TNFR2.subset
prediction.score.GPR183.subset
prediction.score.VIM.subset
prediction.score.HSP.subset
prediction.score.IL7R.subset
Identity
L
GNG4
TNFR2
GPR183
VIM
HSP
IL7R

**Supplemental Figure 3.**

**A**. Reclustered CD4+ T cell UMAPs from ten HBV CNBx participants, colored by identified CD4+ T cell clusters.

**B**. HBV-specific T cell clonotypes from literature overlaid onto the Tfh cell cluster.

**C**. Distribution of six Tfh subsets across participants.

**D**. Differential gene expression for each Tfh subset compared against all other Tfh clusters.

**E**. Shannon's entropy of T cell receptor (TCR) clones within each Tfh subset stratified by participant.

**F**. Pseudotime trajectory map of Tfh development via Slingshot analysis overlaid on the UMAP space.

**G**. Developmental distribution and density of individual Tfh subsets along the pseudotime trajectory line (0 = Root).

**H**. Top 10 specific regulons ranked by Regulon Specificity Score (RSS) for Tfh subsets.

**I**. Unsupervised hierarchical clustering of the top 10 specific regulons and their respective RSS scores across Tfh subsets.

**J**. Hematoxylin and eosin (H&E) stained section of tonsil used for spatial transcriptomics.

**K**. Distribution of transfer scores for each Tfh subset across spatial tissue spots.

**L**. Predicted transfer score for each Tfh subset overlaid onto the tissue architecture. White arrows indicate bins with high enrichment.

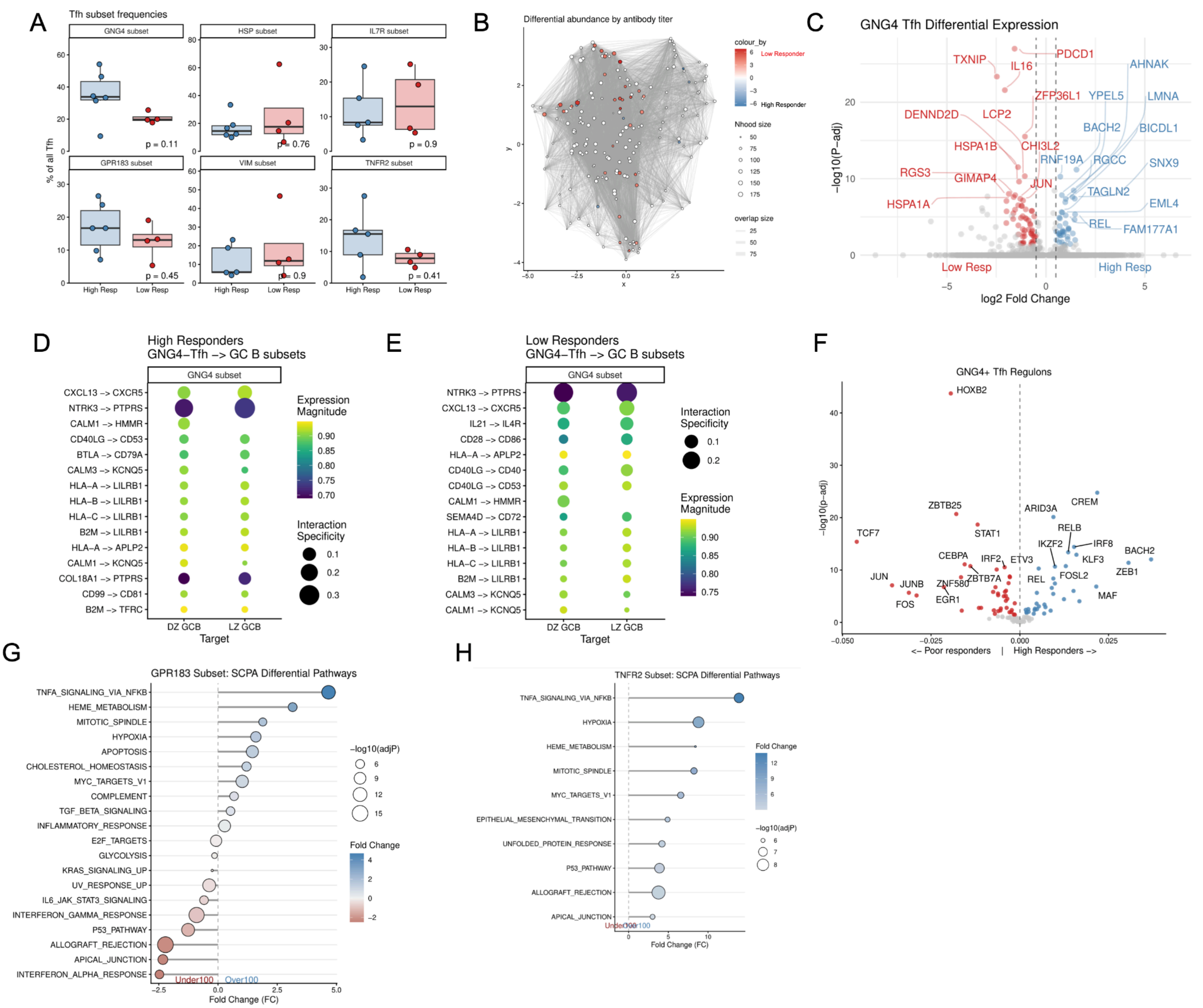


**Supplemental Figure 4.**

**A**. Frequency of six Tfh subsets (as percentage of total Tfh cells) between High Responders and Low Responders. P-values by Wilcoxon test.

**B**. Neighborhood network graph of differential cell abundance associated with antibody titers. Nodes represent cell neighborhoods, colored by log fold change (red indicates enrichment in Low Responders and blue indicates enrichment in High Responders).

**C**. GNG4+Tfh differential expression between Low Responders (red) and High Responders (blue).

**D-E.** Predicted ligand-receptor interactions originating from the GNG4+Tfh subset targeting Dark Zone (DZ) and Light Zone (LZ) Germinal Center (GC) B cells in (**D**) High Responders and (**E**) Low Responders. Circle size indicates NATMI-based interaction specificity, and the color gradient represents expression magnitude.

**F**. Differential transcription factor regulon activity between Low Responders (red) and High Responders (blue).

**G-H**. Single-Cell Pathway Analysis (SCPA) differential MSigDB Hallmark pathways for the (**G**) GPR183 subset and (**H**) TNFR2 subset. Pathways are ranked by fold change.
